# Variant Classification Using Proteomics-Informed Large Language Models Increases Power of Rare Variant Association Studies and Enhances Target Discovery

**DOI:** 10.1101/2025.04.28.650692

**Authors:** Christopher E. Gillies, Joelle Mbatchou, Lukas Habegger, Michael D. Kessler, Suying Bao, Suganthi Balasubramanian, Olivier Delaneau, Jack A. Kosmicki, The Regeneron Genetics Center, Cristen J. Willer, Hyun Min Kang, Aris Baras, Jeffrey G Reid, Jonathan Marchini, Gonçalo R. Abecasis, Maya Ghoussaini

## Abstract

Rare variant association analysis, which assesses the aggregate effect of rare damaging variants within a gene, is a powerful strategy for advancing knowledge of human biology. Numerous models have been proposed to identify damaging coding variants, with the most recent ones employing deep learning and large language models (LLM) to predict the impact of changes in coding sequences. Here, we use newly available proteomics data on 2,898 proteins across 46, 665 individuals to evaluate and refine LLM predictors of damaging variants. Using one of these refined models, we evaluate association between rare damaging variants and human phenotypes at 241 positive control gene-trait pairs. Among these gene-trait pairs, our proteomics-guided model outperforms an ensemble of conventional approaches including PolyPhen2, Mutation Taster, SIFT, and LRT, as well as newer machine learning approaches for identifying damaging missense variants, such as CADD, ESM-1v, ESM-1b and AlphaMissense. When attempting to recover known associations by correctly separating damaging singleton missense variants from other singleton variants, our approach recapitulates 36.5% of gene-trait pairs with known associations, exceeding all the alternatives we considered. Furthermore, when we apply our model to 10 exemplary traits from the UK Biobank, we identify 177 gene-trait associations – again exceeding all other approaches. Our results demonstrate that summary statistics from large-scale human proteomics data enable evaluation and refinement of coding variant classification LLMs, improving discovery potential in human genetic studies.

## Introduction

As sequencing efforts have scaled to hundreds of thousands of individuals, rare-variant focused genetic analyses have helped inform human biology^1–6^. These studies use naturally occurring variation to understand the consequences of disrupting the function of each gene. For a small minority of genes, protein truncation and frameshift variants known as putative loss-of-function (pLoF) variants can be observed in sufficient numbers to enable genetic association studies. For most genes, these pLoF variants are too rare, and genetic association studies must aggregate information across rare missense variants of uncertain functional impact^1,7–9^ to have sufficient power to discover new targets. Effectively aggregating evidence across variants relies on distinguishing likely deleterious or damaging variants from other missense variants, and many approaches have been proposed for this task. One popular strategy^1,3,10,11^ involves using an ensemble of expertly tuned functional prediction scores (see methods)^12–15^. More recently, large language models have demonstrated promise at distinguishing between deleterious and benign coding variants^16–19^. While LLMs have been shown to improve variant classification^17^, it is unclear if this translates to increased power in rare variant association studies.

Here, we use proteomic data collected across 46,665 UK Biobank participants to refine LLM models for protein sequences^20–22^. Proteomics can be utilized to establish connections between genetic variants, protein abundance, and diseases^20–27^, thereby providing valuable insights into disease mechanisms and potential therapeutic targets. While proteomic data is expected to measure protein abundance, it also captures protein structure alterations. For example, previous studies suggest both that (1) 86% of missense cis protein quantitative trait loci’s (pQTLs) variants are associated with decreased protein readouts and that (2) missense cis-pQTLs are enriched for predicted damaging variants^21,28^. We thus hypothesized that missense variants that lead to alterations in protein structure would correspond to both altered protein function and altered proteomic assay readouts^29^ (**Figure 1A**). Parallel work has shown the value of fine tuning LLMs (including protein language models) for specific tasks^30,31^. Therefore, we further hypothesized that measured alterations in proteomic readouts could be used to supervise LLM predictors leading to improved rare variant classification. To assess our hypothesis, we developed a proteomic-refined LLM (**Figure 1B, C**) and compared it with other strategies for identifying damaging missense variants, including an ensemble of conventional approaches, CADD, and LLMs such as ESM-1v^16^, ESM-1b^17,32^ and AlphaMissense^18^ (**Supplemental Figure 1**). We assess our refined LLM in four ways: (1) by assessing correlation of proteomic effect size of genetic variants with predicted deleteriousness scores, (2) by evaluating association between damaging missense variants (identified by our LLM or other classifiers) and specific traits at 241 gene-trait pairs where an association between loss-of-function and the trait is known^1^; (3) by applying our method to gene-level association analysis of ten example traits from UK Biobank; and (4) comparing scores from our proteomics guided LLM for benign and pathogenic clinical variants from ProteinGym^33^ (a benchmark for comparing predicted missense variant effects).

**Figure 1.**
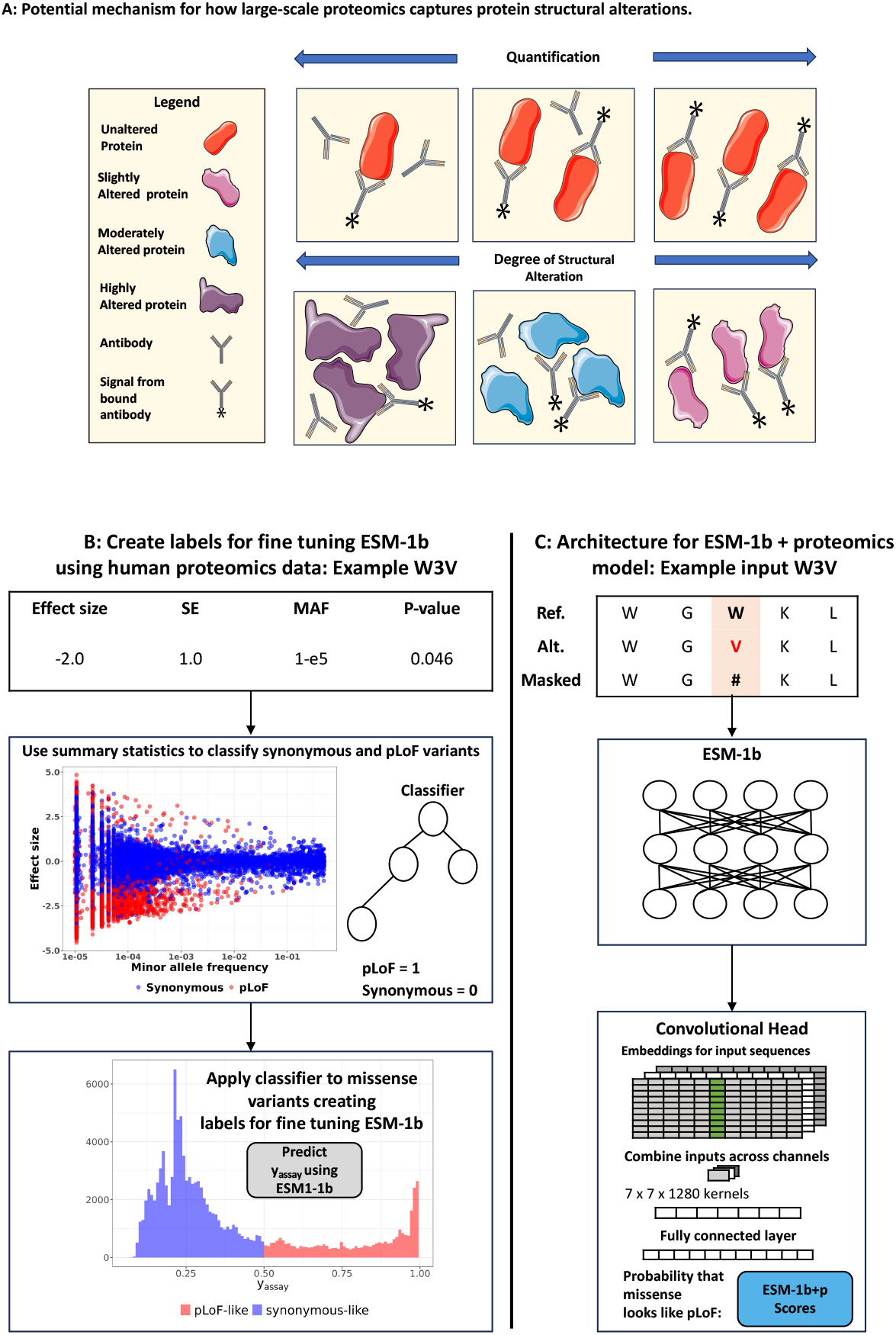
Framework for Finetuning a Large Language Model Using Proteomics Data. **A**. Proteomic assay readouts are affected by coding variants even when abundance remains the same. In the top row, unaltered proteins show increased quantification due to higher abundance, while in the bottom row, poor antibody binding causes diminished quantification, ranging from high to slight protein alterations. Structural changes, other protein modifications (e.g., post-translation modifications) or technical artifacts (slight alterations to antibody binding sites) may result in lower assay readout. Please note to simplify the diagram, we did not include the dual antibodies for Olink’s proximity extension assay. In this figure, clip art was used from Servier Medical Art by Servier, licensed under a Creative Commons Attribution 4.0 International License (https://creativecommons.org/licenses/by/4.0/). Available at: https://smart.servier.com/image-set-download/. **B**. Label Creation for Fine-Tuning ESM-1b. We build a classifier for synonymous or pLoF variants using summary statistics from pQTL analysis. We then apply the classifier to missense variants to generate continuous labels (y_assay_) for fine-tuning ESM-1b. **C**. ESM-1b+Proteomics Model Architecture. We modify the ESM-1b model by adding a convolutional neural network head. We input reference, alternative, and masked protein sequences, and use latent features with the convolutional head to predict y_assay_ values.

## Results

### Protein assay readouts correlate with missense variant deleteriousness

Proteomic assays used the Olink Explore 3072 platform on 46,665 UK Biobank participants of European ancestry who had available exome sequence data. These proteins are distributed between Olink Panel I (n=1,454) and Olink Panel II (n=1,444), which we used for training and validation purposes, respectively (**Supplemental Table 1**). For the 2,898 proteins measured, 560,295 coding variants were observed among assayed individuals. In our analysis of *cis*-coding variants, we found 11,682 variants associated with proteomic assay results (p < 1×10^−8^, see methods^21^) across 1,924 proteins (1,168 Panel I, 756 Panel II). Our training and validation sets included only proteins that had at least one *cis-*coding variant significantly associated with assay readouts.

We first explored evidence for our hypothesis that proteomic assay results were correlated with variant deleteriousness. **Figure 2A** shows that when we group all coding variants (regardless of significance) into three categories (synonymous, missense, and putative loss of function), the estimated impact of each variant on assay readout correlates strongly with the variant type (as expected^21^). Briefly, synonymous variants had no impact on assay readouts (median effect size β = -0.01 +-0.81 standard deviation units, SD) whereas pLoF variants had the largest magnitude impact on assay readouts (median β = -1.40 +-1.35 SD), corresponding to an estimated reduction of 40% in estimated binding (on raw scale). The impact of missense variants was intermediate between these two (median β = -0.20 +-1.05 SD). More remarkably, **Figure 2B** shows that when we group missense variants (regardless of *cis-*pQTL significance) according to deleteriousness as quantified by an ensemble of classifiers (number of deleterious predictions from PolyPhen^12^, HVAR/HDIV, MutationTaster^13^, SIFT^14^, and LRT^15^), the smallest effects on protein assay results are seen for variants predicted to be benign (median β = -0.11 +-0.92 SD) while the largest effects are seen for those predicted to be deleterious (median β = -0.5 +-1.23 SD). **Figure 2C** further shows that proteomic *cis*-pQTL effect sizes for ClinVar benign variants (median β = -0.09 +-0.17 SD) and ClinVar pathogenic variants (median β = -1.22 +-1.16 SD) are also significantly different (p-value = 6.72×10^−48^) effect size. ClinVar pathogenic and benign classifications are used to distinguish likely disease-causing variants from other variants within human disease genes. Not only did we find significantly different association with proteomic assay readouts between ClinVar pathogenic and benign variants, but we also found that proteomic assays results could be used to classify pathogenic and benign variants significantly better (AUROC = 0.76) than a random classifier (AUROC = 0.50) or a classifier using minor allele frequency alone (AUROC = 0.65), consistent with a results from a previous study^21^.These results suggest that effect size from pQTL analysis is a more effective classifier of pathogenic and benign variants than minor allele frequency alone, as evidenced by a higher AUROC score. Together, the observations in **Figure 2** support our hypothesis protein that proteomic assays capture not only protein abundance but also provide a quantifiable measure of missense variants deleteriousness. Most likely, this is because these proteomic assay readouts are sensitive to changes in protein conformation and deleterious variants will often alter protein tertiary structure.

**Figure 2.**
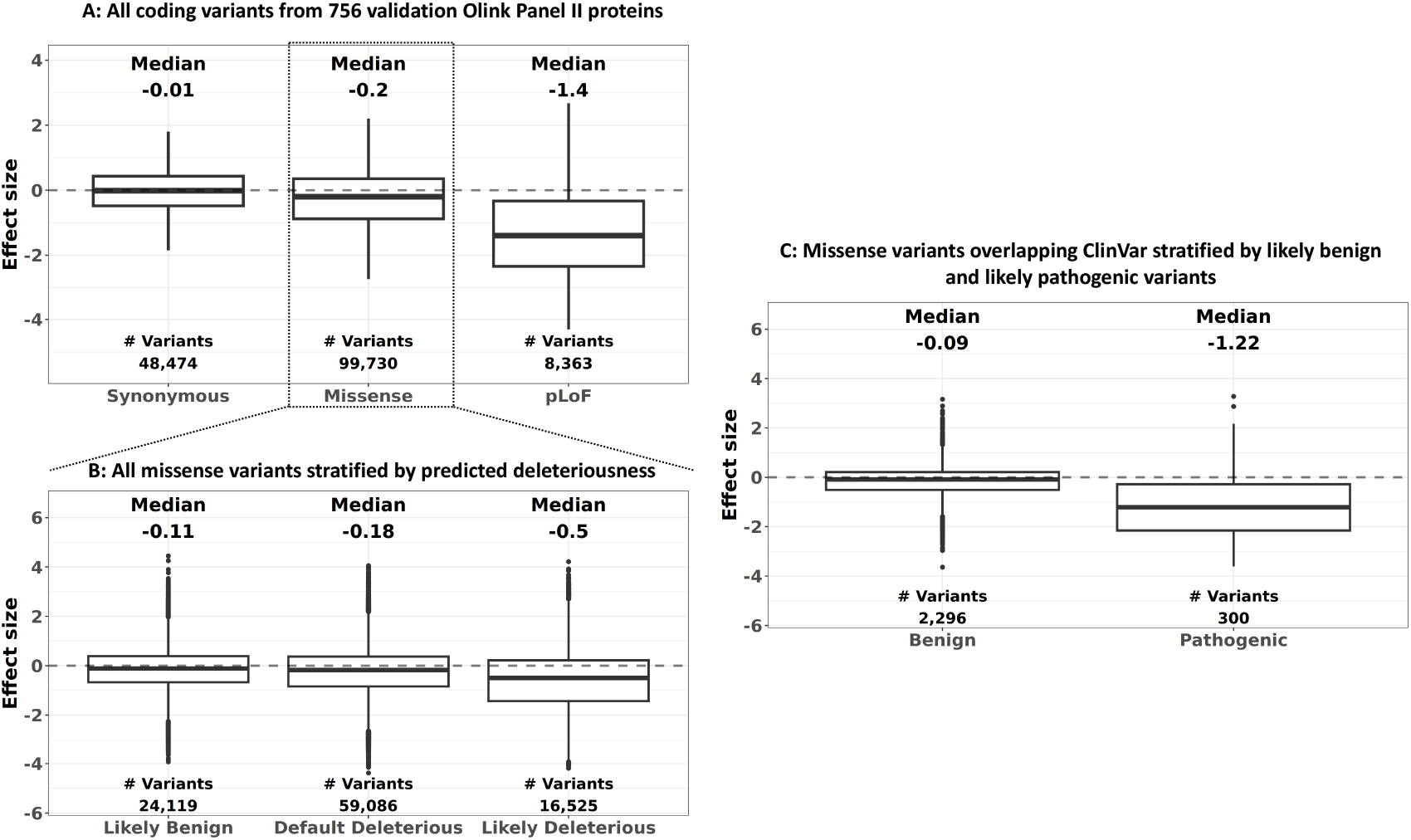
Variant classifica-on on valida-on Olink proteomic data. We plot the distribution of effect size from pQTL analysis classifying all coding variants from 756 Olink Panel II proteins with a cis-coding variant p < 1×10^−8^. **A**. Effect Size of All Coding Variants by Predicted Type on 756 proteins with a significant coding variant. Synonymous variants have a median effect size near zero, pLoFs show large negative effect sizes, and missense variants fall in the middle. **B**. All missense variants stratified by predicted deleteriousness. We used a conventional ensemble that quantify deleteriousness. The ensemble has five functional scores (PolyPhen-2 HDIV/HVAR, LRT, Mutation Taster and SIFT), where Likely Benign = 0 deleterious predictions from all five scores, Default deleterious = 1 to 4 deleterious predictions and Likely Deleterious = 5 deleterious predictions. More deleterious missense variants have larger magnitude effect size. **C**. Missense variants overlapping ClinVar stratified by likely benign and likely pathogenic variants. Only 1-star or higher quality variants were included. We used both benign/likely benign and pathogenic/likely pathogenic variants. The magnitude of effect size for pathogenic variants is significantly larger than benign variants.

### Construction of proteomic-informed LLM utilizing summary statistics from protein quantitative loci analysis

Our next goal was to use proteomic assays results to improve LLM classifiers of variant deleteriousness. We proceeded in two steps. First, we built a classifier that used proteomic assay results to classify missense variants as “pLoF like” (generally, with larger impact on assay readouts and smaller allele frequency) or “synonymous like” (generally, with little impact on assay readouts and relatively higher allele frequencies). In a subsequent step, we used the output of this classifier to refine LLM predictors of deleterious variants.

Thus, in the first step we annotated each variant with allele frequency, proteomics effect size, standard error and p-value (**Supplemental Figure 2)**. Then, we built an ensemble classifier (**Figure 1B, methods**) that used these four features to distinguish benign variants (synonymous variants in Panel I proteins as negative examples) from deleterious variants (pLoF variants in Panel I proteins as positive examples). Once trained, the classifier allowed us to score the deleteriousness of 118,647 observed missense variants with allele frequency <1% observed in Panel I proteins. The output of this classifier was better at distinguishing ClinVar pathogenic and benign variants (AUROC 0.83 in Panel I proteins, 0.79 in Panel II) than classifiers using only effect sizes (AUROC 0.78 in Panel I, 0.76 in Panel II) and those using only allele frequencies (AUROC 0.72 in Panel I, 0.65 in Panel II) (**Supplemental Table 2**).

In the second step, we fine-tuned the ESM-1b proteomics LLM^32^ using results of the classifier for 118,647 missense variants. We call our method ESM-1b+proteomics or ESM-1b+p. The training variants were used as continuous supervised labels, where we added a convolutional neural network head to ESM-1b to pool hidden features from its final transformer layer. We used a small sample of 2,000 random ClinVar pathogenic and benign variants to determine the optimal point for the model to stop training, thereby reducing overfitting to the Olink proteomic platform-specific effects. The optimal point was determined by comparing Spearman correlation of pathogenic and benign variants with the predicted deleteriousness. Our models achieved a Spearman correlation above 0.70 during training compared to 0.67 for standard ESM-1b and 0.62 for ESM-1v on these 2,000 variants. This ESM-1b+p model allowed us to generate proteomic informed missense deleteriousness scores for 73,490,006 potential missense variants across 19,596 proteins in the human genome.

### Missense variant predictions from proteomics-guided LLM are more correlated with validation proteomic assay readouts than all other tested approaches

We performed several experiments to evaluate the performance of our refined LLM compared to other approaches using the 756 Panel II validation proteins. First, for a sanity check, we compared Panel II proteomic assay readouts for 99,730 missense variants not included in training with our ESM-1b+p score and with several other methods for predicting variant deleteriousness. In each evaluation, we grouped predictions for each method into three categories (Likely Benign, Default, Likely Deleterious) matching the category sizes to those from the 5-method conventional ensemble (**Figure 2B**). We compared our LLM with five other approaches ESM-1b, ESM-1v, AlphaMissense, CADD and the 5-method conventional ensemble (the ensemble scores the number of deleterious predictions from PolyPhen^12^, HVAR/HDIV, MutationTaster^13^, SIFT^14^, and LRT^15^ in a scale of 0 – 5 and assigns variants with a score of zero as Likely Benign and those with a score of 5 as Likely Deleterious). **Figure 3A** shows that likely deleterious variants based on the protein-guided LLM had the highest impact on protein assay results among the approaches compared.

**Figure 3.**
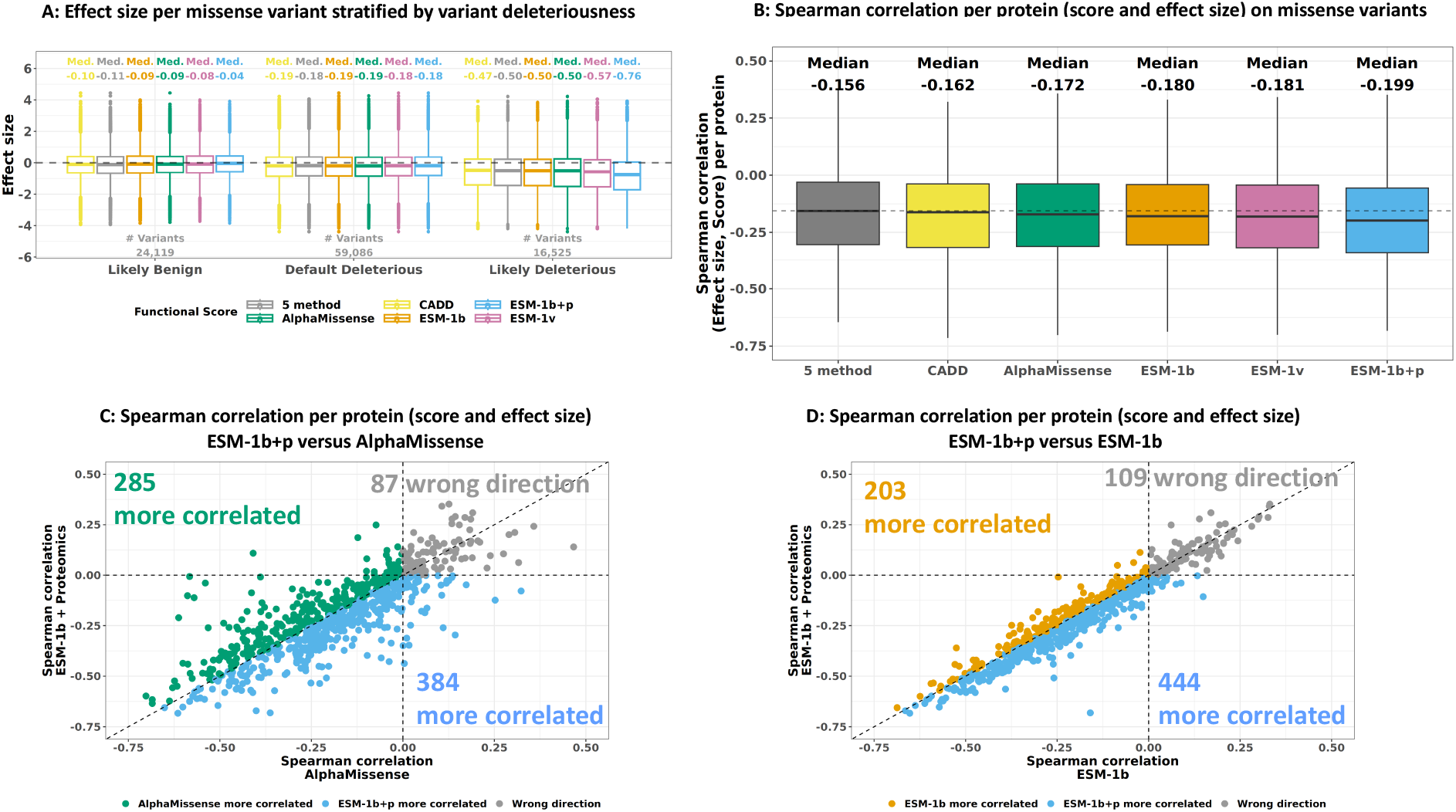
Correlation Analysis of ESM-1b + Proteomics Predictions with Effect Size from Proteomics Analysis Compared to State-of-the-Art Methods. **A**. Effect size per missense variant stratified by variant deleteriousness on 756 validation proteins with a significant coding variant (p < 1×10^−8^). The conventional approach has five functional scores (PolyPhen-2 HDIV/HVAR, LRT, Mutation Taster and SIFT), where Likely Benign = 0 deleterious predictions from all five scores, Default deleterious = 1 to 4 deleterious predictions and Likely Deleterious = 5 deleterious predictions. We matched the category size (number of variants) of all other approaches by ranking missense per method from least to most deleterious. **B**. Spearman Correlation per Protein. ESM-1b + proteomics achieves the largest magnitude median Spearman correlation between predicted deleteriousness and effect size for missense variants using held-out validation proteins. **C**. Comparison with AlphaMissense. Each point represents a protein and corresponds to the Spearman correlation between AlphaMissense or ESM-1b+proteomics’s predictions with effect size from proteomic analysis. Among 669 genes with negative correlations from either method, ESM-1b+proteomics shows stronger correlation with effect size 384 times (blue), compared to 285 times for AlphaMissense (green). **D**. Comparison with Standard ESM-1b. Among 647 genes with negative correlations from either method, ESM-1b + proteomics shows stronger correlation with effect size 444 times (blue), compared to 203 times for standard ESM-1b (orange).

In principle, a method for predicting variant deleteriousness might work well by distinguishing deleterious variants from benign variants *within a gene*, or by distinguishing genes that are tolerant of variation and mostly contain benign variants from those that are not and might be enriched for deleterious variants. To explore the value of our *within gene* predictions, we evaluated the correlation between ESM1b+p scores and proteomic assay readouts within the 756 Panel II genes. **Figure 3B** shows a box plot of the resulting within gene correlation coefficients, where larger magnitude negative values are better. For this metric, the proteomics-guided LLM outperformed the other methods with a median Spearman correlation of -0.199, followed by ESM-1b with -0.180 and AlphaMissense with -0.172. Among proteins with a negative correlation for either ESM-1b+proteomics or AlphaMissense, ESM-1b+proteomics had a stronger correlation 57.4% of the time (p=7.4×10^−5^; **Figure 3C**). For standard ESM-1b, ESM-1b+proteomics had a stronger correlation 68.6% of the time (p=6.6×10^−22^; **Figure 3D**).

Finally, we examined if the within-protein Spearman correlation extended across all missense variants and validation proteins. We compared the overall Spearman correlation between effect size and predicted scores across all missense variants and validation proteins with scores for all methods **(Supplemental Table 3)**. ESM-1b+p had a Spearman correlation of -0.212 compared to -0.160 for ESM-1v, -0.141 for standard ESM-1b, and -0.137 for AlphaMissense.

Note that, while encouraging, these initial observations are somewhat expected – since protein assay results were used to train our ESM1b+p, whereas the other models were trained using other cues about variant deleteriousness.

### Proteomics-guided LLM recapitulates the most pLoF-defined gold standard gene-trait pairs using only missense variants in burden tests

While our results so far show that our proteomic-refined model outperformed other methods when predicting proteomic assay readouts, our primary goal was to determine whether our proteomic-guided model could improve power for genetic association studies. To evaluate this, we selected 241 positive control autosomal gene-trait pairs where pLoF burden tests had previously established associations with a health-related trait^1^. These 241 genes did not overlap Olink Panel I proteins **(Supplemental Table 4)** used for training. We reasoned that it should be possible to recapitulate some of these pLoF burden signals using deleterious missense variants and that improved classification of missense variants would increase the proportion of signals that could be recapitulated. Thus, we classified singleton missense variants in the full UK Biobank cohort with our ESM1b+p classifier and a variety of alternative variant classification strategies and then attempted to recapitulate each of the 241 association signals. This benchmark is quite challenging since even a perfect classifier may not be able to reconstruct a pLoF association using singleton missense variants alone.

In the context of genetic association studies, the optimal balance between a small number of very deleterious variants and a larger number of more modestly deleterious variants is not obvious. To facilitate this balancing act, we ranked variants from most deleterious to least deleterious using each method and then used a method specific grid search to find the optimal method-specific split of likely deleterious variants to include in a genetic association burden test (see methods for details). When using singleton missense variants, we found that ESM-1b+proteomics recapitulated the 88 gene-trait associations with p < 0.01 (**Figure 4A**). This outperformed all alternatives we considered, including using all singleton missense variants (63), using the 5 method ensemble (69), using ESM-1v (73), using the original ESM-1b (83), and using AlphaMissense (87). The relative advantage of ESM-1b+proteomics over other methods increased with more stringent p-value thresholds (**Supplemental Figure 3**). For example, using p-value < 2.1×10^−4^ and singleton missense variants, ESM-1b+proteomics recapitulated 56 gene-trait pairing compared to 51 for AlphaMissense and 50 standard ESM-1b.

**Figure 4.**
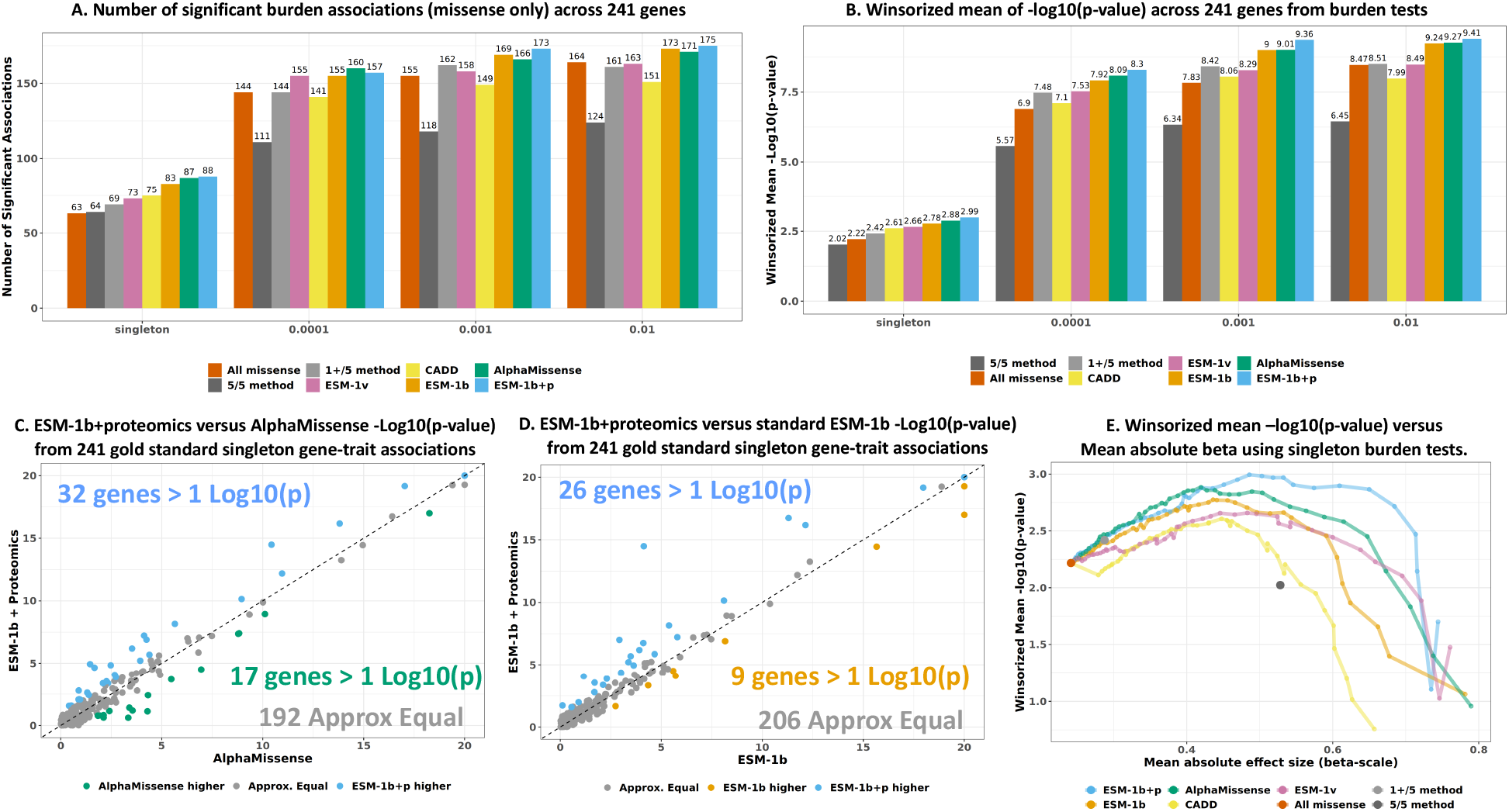
ESM-1b + Proteomics Yields Higher Results on 241 Gold Standard Genes using Burden Tests. **A**. Significant Burden Associations (Missense Only). The number of significant burden associations (p-value < 0.01) across 241 gold standard genes was plotted using only missense variants from different allele frequency bins, where each bin includes all variants up to the specified maximum frequency. Scores were ranked from lowest to highest, and all thresholds were tested with a step size of 0.02. We selected the threshold per method with the highest Winsorized mean -log10(p-value) using singleton missense variants. The same threshold is used for panels A, C, and D. **B**. Winsorized mean of -log10(p-value) across 241 gold standard genes using burden tests. To account for outlier signals, we Winsorized the -log10(p-values) at 20. **C**. ESM-1b + proteomics versus AlphaMissense -log10(p-value) from 241 gold standard singleton gene-trait associations. We indicate the number of times each method showed > 1 order of magnitude improvement. **D**. ESM-1b + Proteomics versus standard ESM-1b -log10(p-value) from 241 gold standard singleton gene-trait associations. **E**. Winsorized mean -log10(p-value) versus mean absolute effect size for all methods. Each point represents a different deleteriousness threshold for each model. For a given threshold, all missense variants with predicted score above the threshold are included in a burden test. All missense, 1+/5 method and 5/5 method only have a single point because there is no threshold to change. From the burden test results across 241 gold standard gene-trait pairs, we can compute the number of significant associations and the mean effect size.

Figure 4B shows that our method increase not only the number of significant gene-trait pairs but also increased the average association signal strength relative to the other methods (measured by the mean -log10(p-value) after winsorization to reduce the impact of outliers). Specifically, ESM1b+p increased mean -log10(p-value) by 7.5% relative to standard ESM-1b and 3.8% relative to AlphaMissense (**Figure 4B**). Using median -log10(p-value) shows a similar trend as Winsorized mean -log10(p-value) (**Supplemental Figure 4**). As a sensitivity analysis, we expanded our evaluation to higher frequency missense variants and found similar patterns as in the analysis restricted to singletons (**Figure 4A,B**). Relative to AlphaMissense, ESM-1b+p improved 32 associations by >1 order of magnitude in p-value compared to 17 improved associations for AlphaMissense, the next best method (**Figure 4C**). Similarly, our model notably boosted 26 associations compared to 9 for standard ESM-1b (p-value=0.003) (**Figure 4D**).

Finally, we investigated whether our proteomic-guided LLM increased gene-level effect size estimates in burden testing. We hypothesized that a larger effect size would indicate more deleterious variants. To test this, we computed the mean absolute effect size across 241 gene-traits for each deleteriousness threshold. We then plotted the mean absolute effect size against the Winsorized mean -log10(p-value) and the number of significant associations **(Supplemental Figure 5)**. We found that ESM-1b+proteomics exhibited noticeably larger effect sizes and mean - log10(p-values) across most deleteriousness thresholds **(Figure 4E)**.

**Figure 5.**
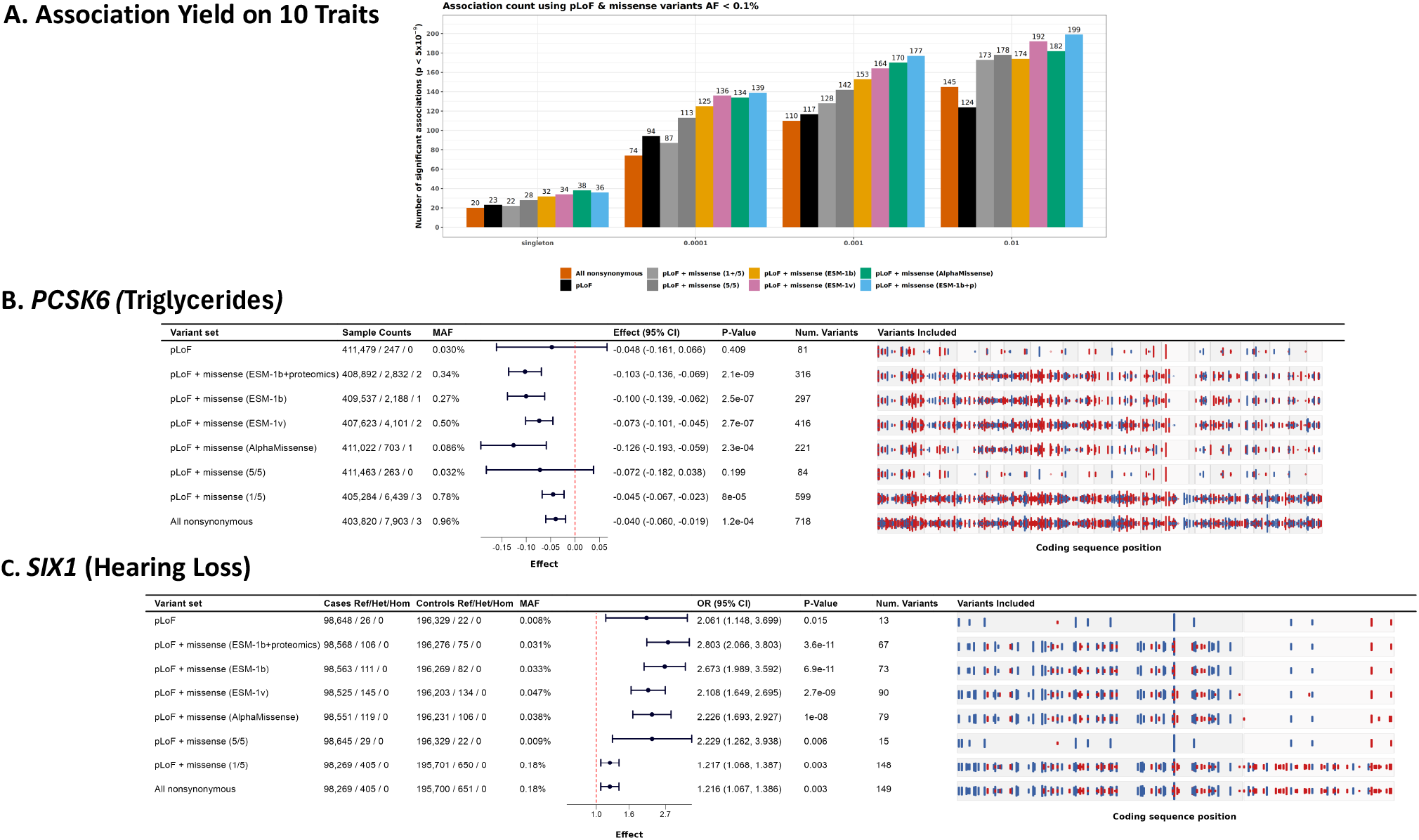
Two Gene-Trait Associations Identified with ESM-1b + Proteomics but Not Standard ESM-1b and AlphaMissense. **A**. Number of significant associations p-value < 5×10^−9^ using burden tests across 10 example UK Biobank traits. **B**. *PCSK6* with triglyceride-levels association. This example shows summary statistics and p-values for burden codings included in the gene-based testing framework (AF < 0.1%). The associations involve pLoF and missense variants with an allele frequency of less than 0.1%. Missense variants were included if they were among the top 22% most deleterious, as determined by AlphaMissense, ESM-1v, ESM-1b, and ESM-1b+proteomics. This panel shows summary statistics and p-values for burden codings included in the gene-based testing framework (AF < 0.1%). The “Sample Counts” column shows genotype counts for homozygous reference, heterozygotes and homozygous alterative counts respectively. The “Variants Included” column displays each variant as a vertical line, with the height corresponding to the magnitude of Z-statistics. Red lines indicate variants with a negative effect size, and blue lines indicate variants with a positive effect size. Exons are shown with different background shades. **C**. *SIX1* association with Hearing Loss.

### Proteomics-guided LLM maximizes yield of associations for ten example UKB traits

We next applied our proteomics-guided model to ten example traits from the UK Biobank (birth weight, body mass index, creatinine, cystatin C, HDL cholesterol, hearing loss, standing height, LDL cholesterol, mean reticulocyte volume, and triglycerides) to contrast association yield with the other approaches for selecting missense variants to include in gene burden tests. To do this evaluation, missense and pLoF variants are combined in groups according to allele frequency (ranging from singletons, 1×10^−5^, 1×10^−4^, 1×10^−3^ to 0.01) and deleteriousness (pLoF, pLoF + deleterious missense, pLoF + default deleterious, pLoF + all missense). As a baseline, deleterious missense variants were defined by all 5 methods in the conventional ensemble predicting the variant to be deleterious, whereas default deleterious requires at least one deleterious prediction. For each LLM approach, we defined the top 22% scoring missense variants as deleterious (over all possible single nucleotide missense variants), and computed combined pLoF and deleterious missense burden tests, comparing to the conventional baseline. This percentage was chosen to match the proportion of deleterious missense variants selected by the 5 method ensemble approach.

When using variants with allele frequency <0.1% and a significance threshold of 5×10^−9^, the 5 method ensemble resulted in 142 significant associations compared to 153, 164, 170 and 177 for ESM-1b, ESM-1v, AlphaMissense and ESM-1b+proteomics, respectively (**Figure 5A**). This translates to a 15.7% increase in yield for ESM-1b+proteomics over standard ESM-1b, and 24.6% increase in yield compared to the 5 method ensemble. The improvement of ESM-1b+proteomics was consistent across allele frequencies and p-value thresholds (**Supplemental Figure 6**). Across these traits, we did not observe inflation of test statistics (**Supplemental Table 5**).

Next, we examined the novel association signals found by our LLM relative to the other methods (**Supplemental Table 6, Supplemental Figure 7**). We highlight one of these in **Figure 5B**. This novel association (Effect=-0.103 SD, p-value=2.1×10^−9^) was for proprotein convertase subtilisin/kexin type 6 (*PCSK6)* with triglycerides. *PCSK6* has common variation associated with triglyceride-levels in the GWAS Catalog^34,35^ and knockout of *PCSK6* is associated with salt-sensitive hypertension in mice^36^. The second best approach for deleterious missense variant prediction in *PCSK6* with triglycerides was standard ESM-1b (Effect=-0.100 SD, p-value=2.5×10^−7^).

Finally, we explored associations that were novel to the LLM approaches relative to the conventional approaches (**Supplemental Table 7**). One of these findings is an association between hearing loss and rare deleterious coding variants in *SIX1*, a gene that plays a critical role in inner ear development during embryogenesis **(Figure 5C)**. This gene has also been connected to hearing loss via de novo mutations^37,38^. When considering variants with allele frequency < 0.1% in *SIX1* in a burden coding, the association using nonsynonymous variants was not significant (Odds ratio = 1.216, p-value=0.003). Using pLoF and missense variants predicted deleterious by 5 algorithms was also not significant (Odds ratio = 2.229, p-value=0.006). All the ESM LLM approaches were significant with the strongest being our approach ESM-1b+proteomics (Odds ratio = 2.803, p-value=3.6×10^−11^).

### Predictions from proteomics-guided LLM improve the classification of ClinVar pathogenic variants over unguided baseline LLM

As a final assessment of the value of proteomics-refinement over standard ESM-1b, we investigate its predictions on ClinVar variants from the ProteinGym^33^ benchmark **(Supplemental Table 8)**. To ensure fair evaluation, we removed ClinVar variants used during training for either model fitting or convergence checking. To ensure balance of pathogenic and benign variants per protein, we selected proteins with at least 4 benign and 4 pathogenic variants, similar to a previous study^31^. This filter resulted in a total of 34,531 variants and 760 genes. ESM-1b+proteomics outperformed standard ESM-1b in both the AUROC and AUPRC for all variants, achieving 0.940 versus 0.919 for AUROC and 0.955 versus 0.936 for area under the precision recall curve (AUPRC), respectively. AlphaMissense performed the best on this metric, achieving an AUROC of 0.947 and an AUPRC of 0.958. A similar pattern also occurred when assessing mean and median AUROC per gene **(Supplemental Table 8)**.

### Proteomic refinement improves performance on a variety of ESM models with differing numbers of parameters showing the robustness of the proteomic training labels

Our analyses so far show that ESM-1b+p outperforms the original ESM-1b model at the task of identifying deleterious missense variants for genetic association studies and, also, that ESM-1b+p outperforms all other approaches we considered for this task We also evaluated whether proteomics data could be used to improve other LLM predictors of variant deleteriousness. For this, we trained a model utilizing ESM-1v model embeddings as input features (**Supplemental Figure 8**) and again found that performance improved over standard ESM-1v when classifying missense singleton variants for a genetic association study (83 versus 73 genes recapitulated), although the refined model performance was lower than for ESM-1b+proteomics (83 versus 88 genes recapitulated) (**Supplemental Figure 9**). We applied proteomics-refinement to various ESM-2 models with sizes ranging from 8 million parameters to 150 million parameters using 2,000 ClinVar variants (our early stopping set). Performance of these models improved with proteomic refinement, but models with fewer parameters improved notably more than models with more parameters (**Supplemental Figure 10**). For example, proteomic-refinement improved ESM-2’s AUROC (with 8 million parameters), from 0.653 to 0.841 exceeding the performance of the 150 million parameter version ESM-2. The 150 million parameter model’s AUROC improved from 0.807 to 0.888 exceeding the performance of the 3 billion parameter model which had an AUROC of 0.878.

## Discussion

Here, we highlight four important advances for genetic association studies of rare coding variation. First, we confirm the value of human proteomics data for comparing various algorithms for damaging missense prediction using summary statistics alone. Second, we show that previously described LLMs trained on protein sequences can improve the power of these studies relative to ensembles of conventional models for variant classification. Third, we show that when these models are trained on the results of proteomic association studies using UK Biobank data, their performance further improves. Fourth, our results suggest that there is orthogonality between our proteomics-guided LLM, AlphaMissense and standard ESM-1b suggesting omnibus tests combining all of these may lead to the most genetic associations. All these show the value of large-scale human proteomics data for empowering genomic discovery.

These advantages were evident in our analysis of 241 pLoF-defined positive control gene-trait pairs both in terms of our ability to recapitulate the pLoF defined association signals and in the improvement of burden association yield of ten UKB traits. Furthermore, our LLM could be leveraged in omnibus tests (including SKAT^39,40^ tests) potentially combining other missense variant classification strategies.

In principle, protein sequence LLMs could also be refined with data from sources such as ClinVar^41,42^, HGMD^28,43^ and Deep Mutational Scans^31,44^. However, large-scale human proteomic data is appealing relative to these alternatives because, when combined with DNA sequence data, it annotates large numbers of variants and is expected to be relatively free of bias^31^. We also provided evidence that the proteomic labels are not largely driven by epitope effects given their classification capabilities on ClinVar pathogenic missense variants and correlation with existing deleterious scores, consistent with previous studies^21^. In the future, an abundance of deeply annotated datasets that combine DNA sequence information, proteomics and other omics might provide even broader training sets that can be used to refine LLMs, either individually or in combination with sources such as ClinVar or HGMD.

To assist with the development of future scores and further improvements, we provide our training labels and validation data. Furthermore, we provide precomputed classification (deleterious, default, likely benign) for all missense variants in canonical transcripts from Ensembl 100 and the annotation files for REGENIE so researchers may utilize these predictions to advance discovery using human genetics data.

There are several interesting future directions for refining our models. For example, leveraging *trans* associations could potentially increase the number of variants in the training set and further enhance variant classification. As with other genetic studies, application of our approach to larger and more diverse samples than the current UK Biobank proteomics resource is also expected to improve accuracy and extensibility of our approach. In conclusion, our findings suggest that refining LLMs using proteomics data can assist in differentiating between deleterious and non-deleterious coding variants, hence enhancing the value of current datasets, and streamlining the annotation process for more discoveries. Incorporation of these ‘newly’ classified deleterious variants into burden tests enhances the analysis of genetic variations and may aid in the discovery of novel target-disease associations.

## Methods

### Protein quantitative trail loci (pQTL) and function annotation

We used data from UK Biobank Pharma Proteomics Project^20^ for pQTL analysis and subsequent model training. Starting with the normalized protein expression (NPX) data from the consortium, we performed principal component analysis and adjusted the NPX data by the top 100 principal components. This was done to adjust for confounders and concomitantly increase power for cis-pQTL detection.

We then ran pQTL analysis using REGENIE^45^ for variants that were captured by UK Biobank exome sequencing efforts^1^ in the 46,665 individuals with blood plasma proteomics data as well as TOPMed variants^46,47^. pQTL regression analysis was run adjusting for age, sex, age x sex, age^2^, and four genetic ancestry principal components as covariates. We used VEP^48^ to annotate variants as synonymous, missense, or loss-of-function (stop gained, essential splice site variants and frameshift indels), using Ensembl Release 100^49^ transcript annotations. We defined canonical transcripts similar to other approaches^2^. Specifically, we selected transcripts with “MANE Select” annotations as the canonical transcript set, the current standard for representing biologically relevant transcripts^50^. In the absence of MANE annotation, we used APPRIS^51^ and “Ensembl canonical” annotation tags to define canonical transcripts. We prioritized APPRIS annotation over “Ensembl canonical” because it uses cross-species conservation as a key feature to identify the likely functional transcript. Ensembl canonical transcript annotation is based on the definition provided here http://jan2019.archive.ensembl.org/info/website/glossary.html

Putative loss-of-function (pLoF) variants were defined as those annotated as frameshift, stop_gained, splice donor, and splice acceptor.

### Definitions of positive control gene-trait pairs

We defined a list of gene-trait pairs with an established pLoF burden signal^1^ as our positive control set of genes. Our inclusion criteria for these gene-trait pairs were that there was a burden signal (p < 2.18×10^−11^, see methods^1^) driven by pLoF variants with allele frequency < 0.01. Since a gene can be associated with many traits, we wanted to reduce the chance a single gene would drive our conclusions. To this end, each gene was selected at most one time, focusing on the most significant trait per gene. This resulted in 276 gene-trait pairs. To further reduce bias in our conclusions from proteomic training, we excluded pairs that overlapped the Panel I Olink proteins used for training, and we focused on autosomal genes, resulting in 242 gene-trait pairs. One gene *LARP1* (ENSG00000155506) was neither 5’ nor 3’ complete using the canonical transcript defined above so we removed it leading to 241 gene-traits.

### Proteomic training labels derived from pLoFs and synonymous variants

To refine the LLMs, we needed a target label to perform supervised training for every missense variant included in our learning panel. We considered several options for creating proteomics training labels including the effect size with inverse variance weighting, but found that combining proteomic summary statistics (effect size, allele frequency, p-value and standard error) into a single missense label improved the classification of ClinVar pathogenic variants. Specifically, we created a model (trained on synonymous and pLoF variants) that generates a continuous score between zero and one indicating the degree to which a missense variant is acting like a pLoF or synonymous variant. Since some proteins may have imprecise antibody binding or a large percentage of its readout below the limit of detection, we limited training proteins to those with a cis-pQTL (p < 3.4×10^−11^), where cis was defined as being within 1mb of the start or the end of the gene. We hypothesized that if there is a cis-pQTL for a protein, then the protein is likely to have a reasonably specific antibody binding. This resulted in 1,175 out of 1,454 genes (from Panel I proteins) having at least one associated cis-pQTL and a missing rate across samples less than 10% for training. From this group of proteins, we selected synonymous and pLoF variants and used them as binary labels (0 and 1, respectively) for training in step one of our two step machine learning approach **(Figure 1B)**. We created an ensemble of two models, which were averaged to get a continuous score per missense variant. We used a neural network trained with pytorch (version 1.13 and pytorch-lightning (version 1.8.1]), and a XGBoost (version 1.6.2) model trained using scikit-learn (version 1.02). The structure of the neural network was a standard three layer fully connected network with a hidden size of 16 and GELU^52^ nonlinear activations. Synonymous and pLoF variants used as labeled training data and missense variants used for prediction. The features used for training consisted of p-value, log10 minor allele frequency, effect size, and standard error. We trained the models using all chromosomes except for chromosome 20, which we used to tune hyperparameters optimizing AUROC. To account for the class imbalance (since there are many more synonymous than pLoF variants), we weighted the samples so they would have an equal total weight across classes. The AUROC of the XGBoost classifier was 0.88 for distinguishing between pLoF and synonymous variants on chromosome 20. The feature importance was (using Shapley values^53^) 0.673 for effect size, 0.149 for p-value, 0.095 for minor allele frequency, and 0.083 for standard error. The neural network classifier with the same features had an AUROC of 0.86 on chromosome 20. Averaging the predictions of these models slightly improved AUROC on overlapping ClinVar variants (**Supplemental Table 2**). The predictions generated from this ensemble model were then used as continuous training labels for refining the ESM models. We call these labels y^_assay^. All AUROC p-value calculations were done using DeLong’s method. The p-values from this approach were close to that form 2,000 bootstraps.

### Refining ESM-1b model with proteomic derived labels

We used the transformers library from Hugging Face to import ESM models. We added a convolutional neural network head to ESM-1b. The input to our modified ESM-model was three amino acid sequences: (1) The reference amino acid sequence, (2) the amino acid sequence with the variant amino acid, and (3) the amino acid sequence masked out. On the forward pass, the ESM-1b model was applied to each of these inputs. The last hidden layer was extracted and concatenated into separate channels. We applied a 1×1 convolution kernel to merge the channels into a single matrix. After this, we applied seven kernels of size seven x hidden size to the resulting matrix with adaptive pooling. We used GELU^52^ nonlinear activations, and we included a drop out operator after this layer. This output was passed into a final fully connected layer. We also added a skip connection from the final hidden layer (selecting out the embedding at the amino acid position for the variant) to the fully connected layer. To reduce dimensionality, we applied a 1×1 convolution to merge across channels before passing into the fully connected layer. Next, we added the log probability difference between the alternative amino acid and reference amino acid at the amino acid sequence position and put this in the final fully connected layer. To adjust for protein length, we also added the log10 of the amino acid sequence length. The length adjustment covariate did not significantly alter the results. To update model weights, we used binary cross entropy between y_assay_ (defined in the previous section) and the predicted value from the forward pass called p_assay_. We also added log(p_REF_) from the masked sequence prediction, where p_REF_ is the likelihood of the reference amino acid, similar to other work^54^. We weighted these by a constant α which we set to 0.01.

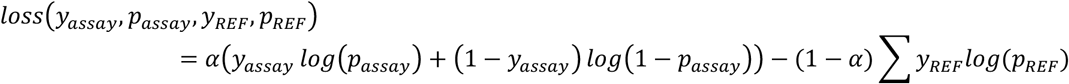

To speed up training, we used low-rank adaptation (LoRA^55^) implemented in the PEFT python package and for training we used the pytorch-lightning. We used 2,000 random benign/likely benign and pathogenic/likely pathogenic variants from ClinVar for early stopping to prevent overfitting. We checked Spearman correlation of the p^assay^ and the binary labels every 20% of an epoch. Once performance stopped improving for one entire epoch, we stopped training taking the best model. To speed up performance, we used a maximum context size of 768. We did not observe a noticeable decrease in performance on the 2,000 ClinVar variants using this context size.

### Augmenting ESM-1v embeddings using supervised learning with proteomics derived labels

The missense scores from the proteomic data are limited to the observed variants and genes, but we can expand to the whole exome by utilizing these missense scores and refining LLMs. The refined LMM can then be applied to all possible missense variants. We first extracted features from the ESM-1v according to the schema shown (**Supplemental Figure 8A**). The approach we took aimed to capture information about gene context and amino acid change for a variant of interest. We designed a 1281×4 matrix per missense variant that captured vectors related to context and sequence change (similar to recently published aproaches^56,57^).

Specifically, we extracted embedding vectors from the final transformer layer of ESM-1v and developed our 1281×4 matrix from a) the average (length 1280) vector of all sequence position vectors spanning a gene b) the (length 1280) vector at a sequence position being considered (which could vary due to a genetic variant) c) another version of the gene context (length 1280) vector, this time including the sequence variant another (length 1280) sequence vector but this time for the variant sequence (as opposed to the reference). We also added a row to this feature matrix (which is how the column vectors go from length 1280 to length 1281) with ESM-1v scores for the variant change in columns 1 and 3 and zeros in columns 2 and 4. The output layer depicted in the schematic in (**Supplemental Figure 8A**). is just for illustrative purposes (i.e. to clarify how the original ESM-1v score was calculated), and is only used in our feature matrix in the final appended row we just described (i.e. the ESM-1v score per variant). As depicted, features are concatenated as columns after sequentially inputting the reference and variant amino acid sequences. The ESM-1v model is publicly available (https://github.com/facebookresearch/esm). Finally, we used the missense scores predicted by our step 1 ensemble model (as continuous training labels on the probability scale) for training/tuning this ESM-1v derived feature matrix. Training was done with a convolutional neural network (CNN) implemented in pytorch (version 1.13) and pytorch-lightning (version 1.8.1), and using the architecture depicted in (**Supplemental Figure 8B**). We used a CNN to take advantage of the input structure that we extract from ESM-1v, as each row corresponds to the same position from the transformer or output layer from ESM-1v. By leveraging a CNN, each row of this matrix could have a kernel applied to it, thus dramatically reducing the parameter space as compared to a standard feed forward network. We used layer normalization^58^ and GELU^52^ nonlinear activations, and we included a drop out operator after the CNN layer. Finally, we used L2 normalization for weight regularization and used binary cross entropy with continuous labeling as our loss function. Some network architectural choices were inspired by GPN^59^.

When training this model, we used 22 of 23 chromosomes, while chromosome 20 was held out of training and used to determine early stopping criteria (i.e. we used it as the “tuning” chromosome). We evaluated training progression using the Spearman correlation statistic, and if this did not increase for 10 epochs, we would stop training and selected the best model for subsequent prediction. The developers of ESM-1v trained five models and the final ESM-1v prediction was as average of the 5 models. We took a similar approach, where we trained a separate CNN per ESM-1v model and took the average of the 5 trained models to make the final prediction for a variant. For genes longer than 1022 amino acids (the maximum context size of ESM-1v), we centered the context at the variant position. We then created predictions for all possible missense variants for canonical transcripts in Ensembl 100.

### Mapping AlphaMissense scores to transcripts used in this study

We downloaded AlphaMissense^18^ scores in March 2024. To map these scores to the transcripts used in our study, we first retrieved the Ensembl gene and transcript identifiers for all canonical transcripts within our gene set. Using the provided AlphaMissense scores, we determined if a prediction was available for the exact transcript used, resulting in 18,308 exact transcript matches. We then attempted to match the remaining genes by gene identifier, resulting in 1,156 genes with non-exact transcript matches. For these genes without exact transcript matches, we assigned the most deleterious score across all transcripts for each possible single nucleotide variant. Finally, we applied a rank score to all potential variants based on the mapped variants, from least to most deleterious. A total of 156 genes could not be matched using this procedure.

### Burden analysis using positive control gene-trait pairs

We used the REGENIE^45^ whole genome regression framework for gene burden association testing. Genetic variants were determined using the 430,998 European exomes from the UK Biobank^1^. As described in previous work^1^, we included variants annotated as ‘start_lost’ and ‘stop_lost’ as missense variants in our missense gene burden codings. We took predicted ESM-1v scores and predicted ESM-1v+proteomics scores per missense variant and rank-transformed these scores (scaled from 0 to 1) over observed variants with AF < 1% among pLoF-associated genes. When doing gene burden analysis, we needed to define a threshold such that all variants above would be considered a variant deleterious (and therefore consequential enough to be included in gene burden codings). We did not want this threshold to lead to erroneous conclusions, so we varied the threshold from zero to 0.98 by 0.02, and then tested gene burden variant inclusion at each threshold.

To evaluate model performance as a function of gene burden association yield, we used multiple criteria. The first criterion we used was the fraction of gold standard gene-trait pairs discovered with nominally significant association signal (p-value < 0.01). We also assessed multiple threshoved to evaluate the robustness of our conclusion. The denominator was calculated as the actual number of gene trait pairs, so that gene-trait pairs missing association p-values (i.e., no variants meeting deleterious threshold) would be penalized. The second criterion we considered was the Winsorized mean -log10(p-value), where we truncated - log10(p-values) at 20, to adjust for outliers. The third criterion we considered was median - log10(p-value). The fourth criterion we used was the mean absolute effect size across all gene-trait pairs. For the head-to-head comparisons, we used a one-sided binomial sign test to see if ESM-1b proteomics had higher numbers of traits with increased association strength.

### Yield analysis on ten UKB traits

To assess the performance of ESM-1b+proteomics compared to the other approaches, we applied combined gene-based testing to ten UKB traits (birth weight, body mass index, creatinine, cystatin C, HDL cholesterol, hearing loss, height, LDL cholesterol, mean reticulocyte volume, and triglycerides). Using canonical Ensembl 100 genes (see above), we generated scores from ESM-1b, ESM-1b + proteomics, AlphaMissense and ESM-1v for all possible missense variants and genes. We rank-transformed these scores to a scale from zero to one, with a higher score indicating a more deleterious variant. We defined the top 22% of scores as deleterious based on all possible missense variants. This percentage matched the proportion of variants with 5 deleterious predictions from the conventional ensemble approach. This approach allows a different fraction of deleterious variants per gene. We carried out gene-based association testing using REGENIE^44^. More precisely, we tested multiple groupings of missense and pLoF variants based on the annotation strategy and alternate allele frequency cutoffs (singletons, 0.01%, 0.1%, 0.5% and 1%), and combined all the tests through a unified testing framework to result in a single p-value for each gene/trait pair^28^. We defined a genome-wide significance threshold of 5×10^−9^ corresponding to a Bonferroni correction for the number of traits, genes and annotation strategies compared.

