## Supplemental Figures for "Variant Classification Using Proteomics-Informed Large Language Models Increases Power of Rare Variant Association Studies and Enhances Target Discovery"

### Supplemental Figures and Tables

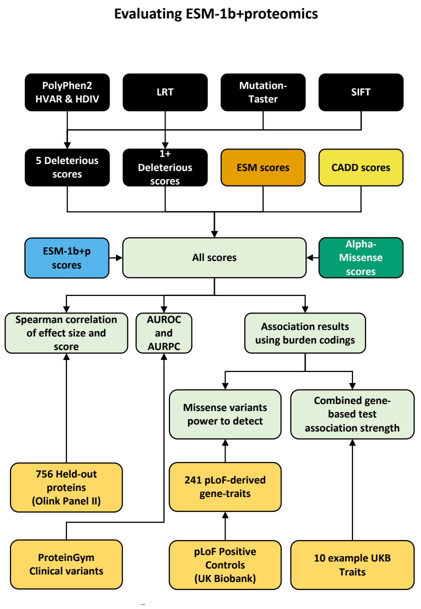


**Supplemental Figure 1.** Evaluation of LLMs' Ability to Discover Rare Damaging Variants and Improve Genetic Discovery. We compare missense scores from conventional ensemble prediction, ESM-1v, ESM-1b, AlphaMissense, and our ESM-1b+proteomics. We evaluate these scores in four ways: (1) Spearman correlation of effect size (beta from pQTL) and deleteriousness scores. (2) Gene codings created from scores' ability to reproduce pLoF-derived positive control gene-trait pairs. (3) Example application across ten UKB traits including: birth weight, body mass index, creatinine, cystatin C, HDL cholesterol, hearing loss, height, LDL cholesterol, mean reticulocyte volume, and triglycerides. (4) Area under the receiver operating characteristic (AUROC) and precision recall curves (AUPRC) on clinical variants from ProteinGym.

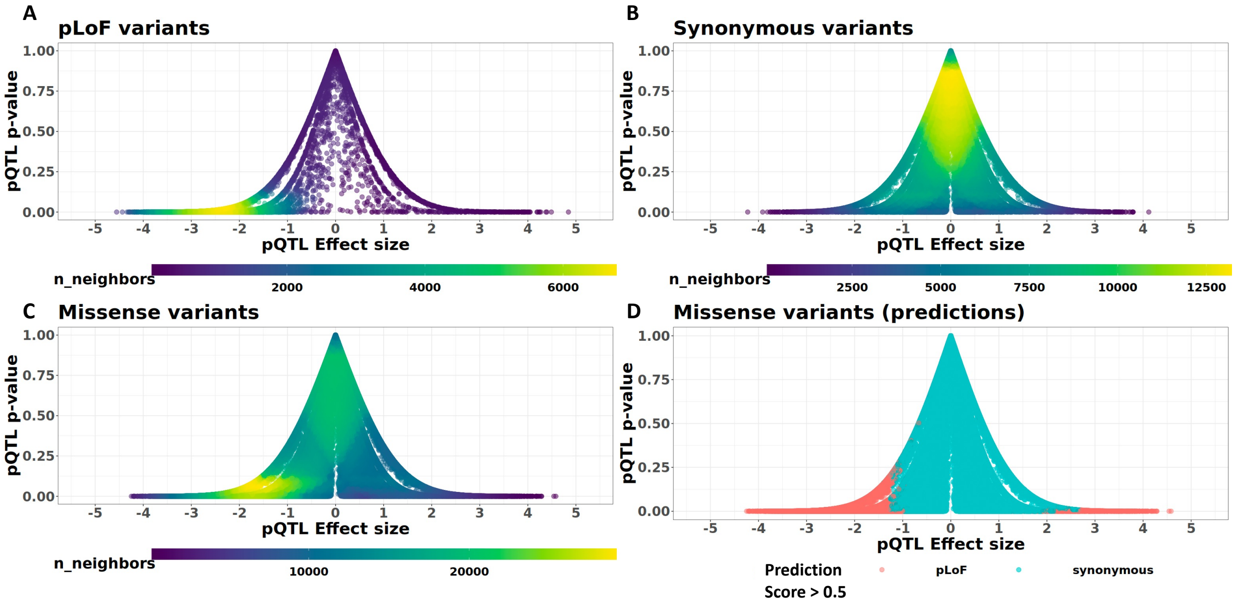


**Supplemental Figure 2: Scatter plot of cis-pQTL effect size and p-value stratified by variant annotation type.**

**A.** Scatter plot of pLoF variants. The x-axis corresponds to the effect size from pQTL and the y-axis corresponds to the p-value. **B.** Synonymous variants. **C.** Missense variants. **D.** Predictions from ensemble model for missense variants using allele frequency, proteomics effect size, standard error and p-value. We trained this ensemble model using pLoF (labeled 1) and synonymous variants (labeled 0) using four features (allele frequency, beta, p-value, and standard error from our pQTL analysis (Olink Panel I proteins). We then applied this model to missense variants to get a probabilistic prediction for how much a missense variant looks like a pLoF. We defined a binary cutoff of 0.5 to color points in the plot for visualization reasons.

| Possible Proteomic Labels For Refining ESM-1b | AUROC | AUPRC |
|---|---|---|
| Proteomic Score: Average XGBoost and Neural Network | 0.829 | 0.543 |
| Proteomic Score: XGBoost Only | 0.828 | 0.531 |
| Proteomic Score: Neural Network Only | 0.824 | 0.531 |
| Effect Size | 0.779 | 0.525 |
| Absolute(Effect Size) | 0.816 | 0.497 |
| Minor allele frequency | 0.703 | 0.371 |
| Z-statistic | 0.71 | 0.221 |
| SE | 0.696 | 0.192 |
| P-value | 0.665 | 0.18 |
| Absolute(Z-statistic) | 0.665 | 0.18 |

**Supplemental Table 2:** An ensemble classifier combining summary statistics from pQTL analysis is most associated with ClinVar variants. We selected 3,484 pathogenic/likely pathogenic (12.3%) and benign/likely benign (87.7%) variants from ClinVar that overlapped variants from the training proteins. We trained two classifiers predicting pLoF and Synonymous from proteomic summary statistics one using a XGBoost classifier and the other using a standard feedforward neural network. We found that averaging the predictions from these classifiers led to the highest area under the receiver operating characteristic and precision recall curves. **AUROC:** Area Under Receiver Operating Characteristic Curve, **AUPRC:** Area Under Precision Recall Curve.

Regeneron - Internal

| | Spearman correlation of Score and Effect Size | Spearman correlation of Score and Absolute Effect Size |
|---|---|---|
| **ESM-1b+proteomics** | -0.212 | 0.209 |
| **ESM-1b** | -0.141 | 0.140 |
| **ESM-1v** | -0.160 | 0.156 |
| **AlphaMissense** | -0.137 | 0.162 |
| **CADD** | -0.131 | 0.148 |
| **5 method** | -0.133 | 0.145 |

**Supplemental Table 3: ESM-1b+Proteomics Correlation with Missense Variant Effect Sizes.** ESM-1b+proteomics shows higher correlation with missense variant effect sizes compared to other deleteriousness scores. We compare ESM-1b+proteomics on 99,730 missense variants scored by all methods across 756 proteins, using Spearman correlation for both effect size and absolute effect size from pQTL analysis.

Regeneron - Internal

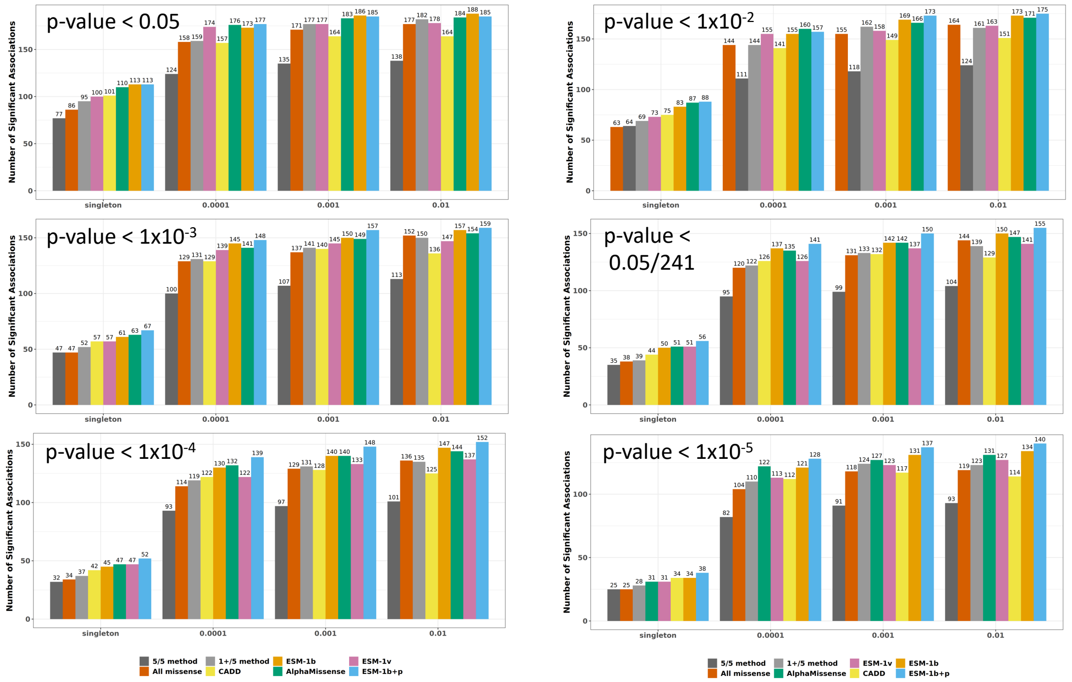


**Supplemental Figure 3: ESM-1b+proteomics improves over standard ESM-1b at more stringent p-value thresholds.** Significant Burden Associations (Missense Only). The number of significant burden associations across 241 gold standard genes (using different p-value thresholds) is plotted using different allele frequency bins, where each bin includes all variants up to the specified maximum frequency. Scores were ranked from lowest to highest, and all thresholds were tested with a step size of 0.02. We selected the threshold per method with the highest Winsorized mean -log10(p-value) using singleton missense variants. The same threshold is used for all analyses in this figure.

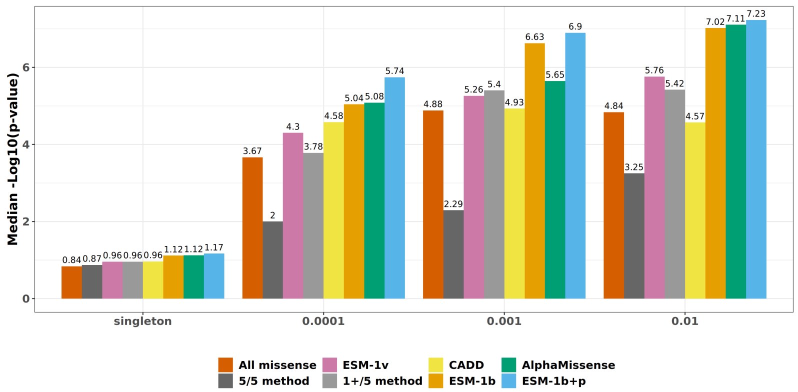


**Supplemental Figure 4: Median –log10(p-value) shows a similar trend as Winsorized mean log10(p-value).** Here we compute the median -log10(p-value) across 241 gold standard gene-trait pairs using burden tests of missense variants.

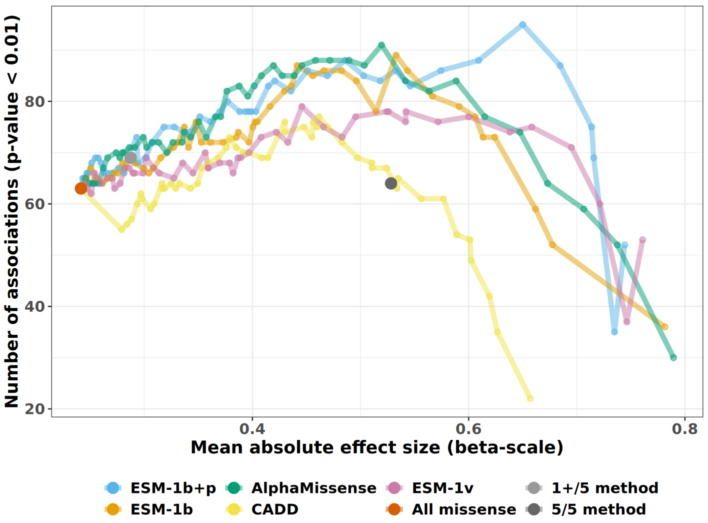


**Supplemental Figure 5: Number of associations (p-value < 0.01) versus mean absolute effect size.** Each point represents a different deleteriousness threshold for each model. For a given threshold, all missense variants with predicted score above the threshold are included in a burden test. All missense, 1+/5 method and 5/5 method only have a single point because there is no threshold to change. From the burden test results across 241 gold standard gene-trait pairs, we computed the number of significant associations and the mean effect size.

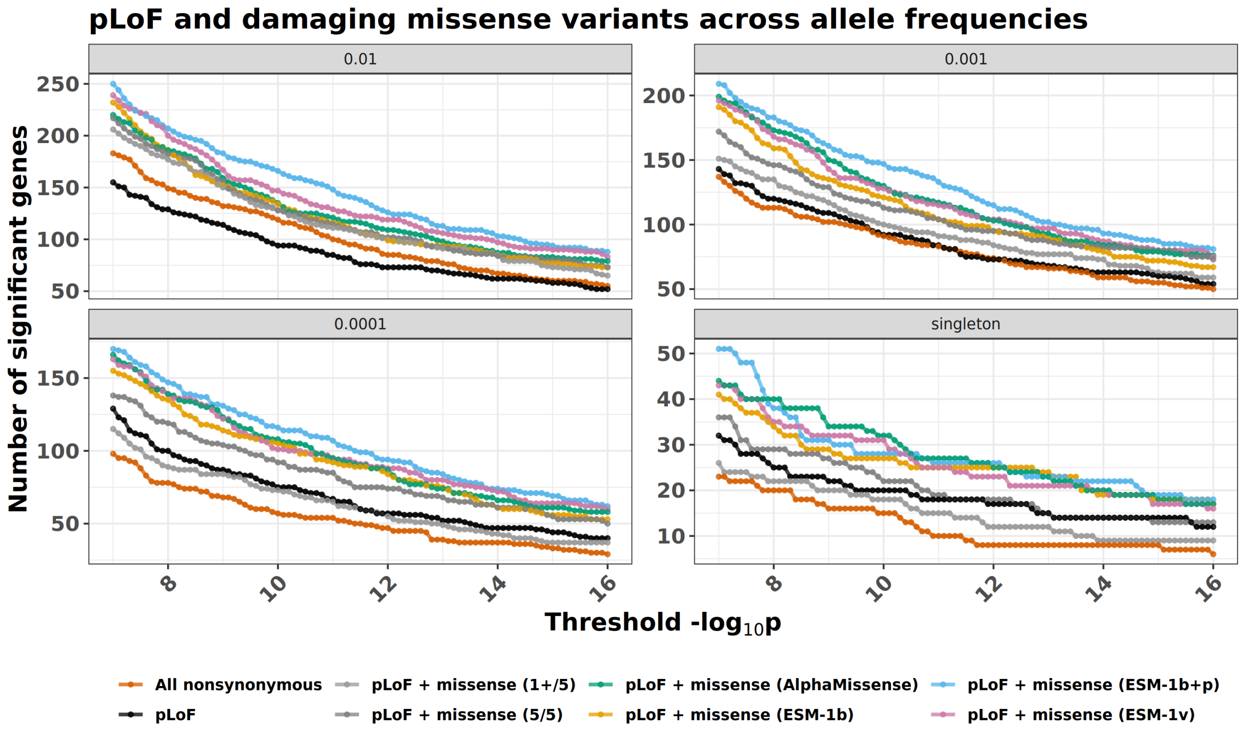


**Supplemental Figure 6: Comparison of Burden Tests Across Allele Frequency and P-Value Thresholds using 10 UKB traits**. We present four panels corresponding to different variant inclusion allele frequencies for the burden tests across all genes from 10 UKB traits. The x-axis in these figures represents the -log10(p-value) threshold used to determine if an association is significant. The y-axis indicates the total number of significant associations. Each color corresponds to a different method.

| Trait | ESM-1b+p | ESM-1b | ESM-1v | AlphaMissense | 1+/5 method | 5/5 method |
|---|---|---|---|---|---|---|
| BMI | 1.144 | 1.116 | 1.133 | 1.15 | 1.129 | 1.122 |
| Mean Reticulocyte Volume | 1.056 | 1.044 | 1.029 | 1.052 | 1.02 | 1.029 |
| Creatinine | 1.093 | 1.063 | 1.065 | 1.083 | 1.083 | 1.068 |
| Cystatin C | 1.126 | 1.064 | 1.1 | 1.083 | 1.067 | 1.08 |
| HDL Cholesterol | 1.044 | 1.059 | 1.046 | 1.075 | 1.029 | 0.995 |
| LDL Cholesterol | 1.031 | 1.025 | 1.025 | 1.037 | 0.99 | 0.993 |
| Triglycerides | 1.036 | 1.034 | 1.05 | 1.028 | 1.037 | 1.041 |
| Standing Height | 1.191 | 1.228 | 1.231 | 1.203 | 1.18 | 1.17 |
| Hearing Loss | 1.05 | 1.044 | 1.07 | 1.057 | 1.038 | 1.05 |

**Supplemental Table 5: ESM-1b+proteomics does not exhibit genomic inflation.**
Here we show the genomic inflation factors for combined pLoF & damaging missense variants using variants with AF < 0.1%. All methods have similar levels of inflation for the 10 UKB traits. For traits like standing height, the inflation across methods is not unexpected due to its polygenic nature.

| Gene | Trait | pLoF | All nonsynonymous | pLoF & missense (1+/5) | pLoF & missense (5/5) | pLoF & AlphaMissense | pLoF & ESM1v | pLoF & ESM1b | pLoF & ESM1bp | Common variant GWAS Catalog? | Training protein? |
|---|---|---|---|---|---|---|---|---|---|---|---|
| *PPARG* | Triglycerides | 2.83E-04 | 3.27E-08 | 3.74E-08 | 7.29E-04 | 5.06E-05 | 3.70E-08 | 2.02E-08 | 1.07E-10 | Yes | No |
| *RRBP1* | LDL Cholesterol | 7.55E-07 | 1.36E-02 | 8.41E-04 | 7.16E-07 | 6.57E-06 | 3.12E-02 | 2.41E-08 | 1.21E-09 | Yes | No |
| *SPHK2* | Mean reticulocyte count | 8.91E-02 | 7.35E-05 | 5.42E-05 | 8.11E-06 | 2.72E-07 | 1.62E-05 | 2.33E-07 | 2.01E-09 | Yes | No |
| *PCSK6* | Triglycerides | 4.09E-01 | 1.19E-04 | 7.97E-05 | 1.99E-01 | 2.30E-04 | 2.66E-07 | 2.51E-07 | 2.10E-09 | Yes | No |

**Supplemental Table 6: Four associations using variants with AF < 0.1% not found with any other approach.** We show the p-value of the association of each approach in each cell. The significance threshold was $p < 5x10^{-9}$. The second to last column corresponds to the presence of a common variant association in the GWAS catalog. The final column indicates if the gene was included in training or not.

### *A. SPHK2* Mean Reticulocyte Volume

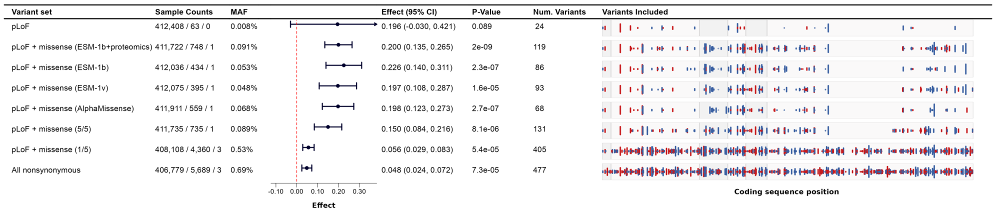


| Variant set | Sample Counts | MAF | Effect (95% CI) | P-Value | Num. Variants |
|---|---|---|---|---|---|
| pLoF | 412,408 / 63 / 0 | 0.008% | 0.196 (-0.030, 0.421) | 0.089 | 24 |
| pLoF + missense (ESM-1b+proteomics) | 411,722 / 748 / 1 | 0.091% | 0.200 (0.135, 0.265) | 2e-09 | 119 |
| pLoF + missense (ESM-1b) | 412,036 / 434 / 1 | 0.053% | 0.226 (0.140, 0.311) | 2.3e-07 | 86 |
| pLoF + missense (ESM-1v) | 412,075 / 395 / 1 | 0.048% | 0.197 (0.108, 0.287) | 1.6e-05 | 93 |
| pLoF + missense (AlphaMissense) | 411,911 / 559 / 1 | 0.068% | 0.198 (0.123, 0.273) | 2.7e-07 | 68 |
| pLoF + missense (5/5) | 411,735 / 735 / 1 | 0.089% | 0.150 (0.084, 0.216) | 8.1e-06 | 131 |
| pLoF + missense (1/5) | 408,108 / 4,360 / 3 | 0.53% | 0.056 (0.029, 0.083) | 5.4e-05 | 405 |
| All nonsynonymous | 406,779 / 5,689 / 3 | 0.69% | 0.048 (0.024, 0.072) | 7.3e-05 | 477 |

### *B. PPARG* Triglycerides

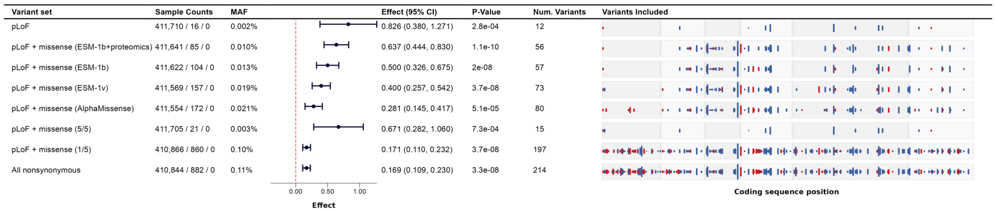


| Variant set | Sample Counts | MAF | Effect (95% CI) | P-Value | Num. Variants |
|---|---|---|---|---|---|
| pLoF | 411,710 / 16 / 0 | 0.002% | 0.826 (0.380, 1.271) | 2.8e-04 | 12 |
| pLoF + missense (ESM-1b+proteomics) | 411,641 / 85 / 0 | 0.010% | 0.637 (0.444, 0.830) | 1.1e-10 | 56 |
| pLoF + missense (ESM-1b) | 411,622 / 104 / 0 | 0.013% | 0.500 (0.326, 0.675) | 2e-08 | 57 |
| pLoF + missense (ESM-1v) | 411,569 / 157 / 0 | 0.019% | 0.400 (0.257, 0.542) | 3.7e-08 | 73 |
| pLoF + missense (AlphaMissense) | 411,554 / 172 / 0 | 0.021% | 0.281 (0.145, 0.417) | 5.1e-05 | 80 |
| pLoF + missense (5/5) | 411,705 / 21 / 0 | 0.003% | 0.671 (0.282, 1.060) | 7.3e-04 | 15 |
| pLoF + missense (1/5) | 410,866 / 860 / 0 | 0.10% | 0.171 (0.110, 0.232) | 3.7e-08 | 197 |
| All nonsynonymous | 410,844 / 882 / 0 | 0.11% | 0.169 (0.109, 0.230) | 3.3e-08 | 214 |

### *C. RRBP1* LDL

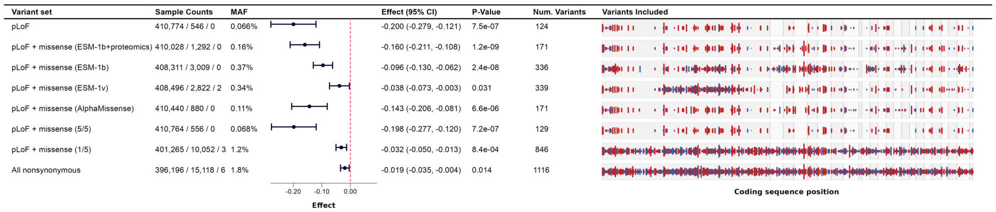


| Variant set | Sample Counts | MAF | Effect (95% CI) | P-Value | Num. Variants |
|---|---|---|---|---|---|
| pLoF | 410,774 / 546 / 0 | 0.066% | -0.200 (-0.279, -0.121) | 7.5e-07 | 124 |
| pLoF + missense (ESM-1b+proteomics) | 410,028 / 1,292 / 0 | 0.16% | -0.160 (-0.211, -0.108) | 1.2e-09 | 171 |
| pLoF + missense (ESM-1b) | 408,311 / 3,009 / 0 | 0.37% | -0.096 (-0.130, -0.062) | 2.4e-08 | 336 |
| pLoF + missense (ESM-1v) | 408,496 / 2,822 / 2 | 0.34% | -0.038 (-0.073, -0.003) | 0.031 | 339 |
| pLoF + missense (AlphaMissense) | 410,440 / 880 / 0 | 0.11% | -0.143 (-0.206, -0.081) | 6.6e-06 | 171 |
| pLoF + missense (5/5) | 410,764 / 556 / 0 | 0.068% | -0.198 (-0.277, -0.120) | 7.2e-07 | 129 |
| pLoF + missense (1/5) | 401,265 / 10,052 / 3 | 1.2% | -0.032 (-0.050, -0.013) | 8.4e-04 | 846 |
| All nonsynonymous | 396,196 / 15,118 / 6 | 1.8% | -0.019 (-0.035, -0.004) | 0.014 | 1116 |

**Supplemental Figure 7: Three additional novel associations.**
Each example shows summary statistics and p-values for burden codings included in the gene-based testing framework (AF < 0.1%). The associations involve pLoF and missense variants with an allele frequency of less than 0.1%. Missense variants were included if they were among the top 22% most deleterious, as determined by AlphaMissense, ESM-1v, ESM-1b, and ESM-1b+proteomics. This panel shows summary statistics and p-values for burden codings included in the gene-based testing framework (AF < 0.1%). The “Sample Counts” column shows genotype counts for homozygous reference, heterozygotes and homozygous alterative counts respectively. The "Variants Included" column displays each variant as a vertical line, with the height corresponding to the magnitude of Z-statistics. Red lines indicate variants with a negative effect size, while blue lines indicate variants with a positive effect size. Exons are shown with different background shades.

Regeneron - Internal

**ESM-1b+proteomics improves performance over ESM-1b on ProteinGym ClinVar variants**

| Model | AUROC all variants | AUPRC all variants | Mean AUROC per gene | Median AUROC per gene |
|---|---|---|---|---|
| AlphaMissense | 0.947 | 0.958 | 0.951 | 0.977 |
| ESM-1b+proteomics | 0.941 | 0.955 | 0.923 | 0.957 |
| ESM-1b | 0.919 | 0.936 | 0.910 | 0.939 |
| EVE | 0.917 | 0.931 | 0.929 | 0.953 |
| ESM-1v | 0.915 | 0.942 | 0.901 | 0.949 |
| CADD | 0.893 | 0.896 | 0.912 | 0.937 |
| 5 method | 0.861 | 0.893 | 0.888 | 0.908 |

**Supplemental Table 8: ESM-1b+proteomics improves performance over ESM-1b on ProteinGym ClinVar variants.** We demonstrate the performance of ESM-1b+proteomics compared to other approaches using ProteinGym ClinVar variants. Overlapping variants included in our early stopping variant set for training ESM-1b+proteomics were removed. We retained proteins with at least 4 benign and 4 pathogenic variants per gene, totaling 35,673 variants and 780 proteins. We compared the AUROC for all variants, as well as the per gene mean and median. Additionally, we report the AUPRC for all variants. ESM-1b+proteomics outperformed standard ESM-1b across all metrics. However, AlphaMissense outperformed ESM-1b+proteomics.

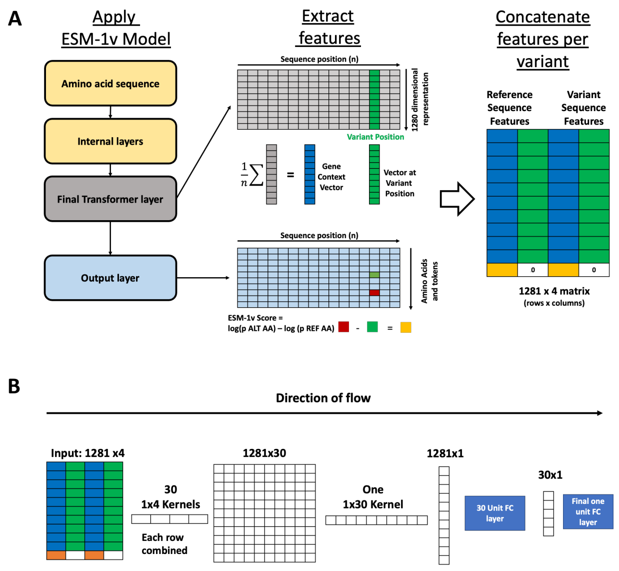


**Supplemental Figure 8: ESM-1v feature extraction and refined model**

**A.** Starting from the left for each variant, we apply the ESM-1v model twice. Once with the reference sequence and once with the variant sequence. We extract features from the final transformer layer and the final output layer. From the transformer layer, we extract a *Gene Context* vector and a *Variant context vector*. The *Gene context vector* is the average of all non-padding over all 1280x1 vectors across the sequence, while the *Variant context vector* is the 1280x1 dimensional vector at the variant position. We also compute the ESM-1v score using the reference and the variant sequence. Finally, the features extracted from the reference and variant sequences are concatenated into a 1281x4 matrix. Each variant will have a 1281x4 matrix as input to downstream machine learning applications. **B.** Convolutional Neural network model for ESM-1v feature refinement for a single variant. Starting from the left we 30 1x4 kernels to each row of the ESM-1v feature matrix. We then apply one 30x1 kernel to summarize the resulting tensor to a 1281x1 vector. Next, 30 fully connected linear units with gaussian error linear unit activation functions are applied to this 1281x1 vector. A final fully connected layer is applied with a sigmoid activation function to give an output that ranges from zero to one, where one is most like a pLoF variant.

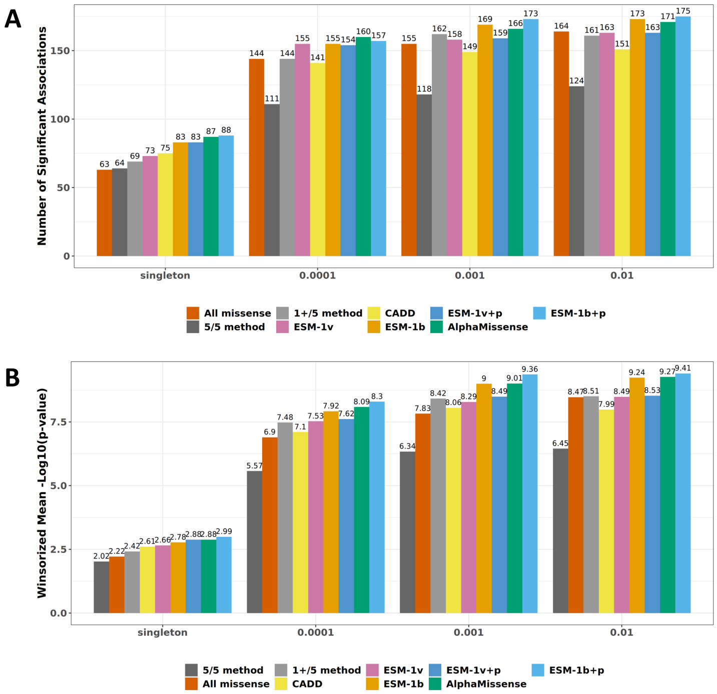


**Supplemental Figure 9: ESM-1v with proteomics fine tuning improves over standard ESM-1v but is worse than ESM-1b + proteomics fine tuning.**
**A.** Significant Burden Associations (Missense Only). The number of significant burden associations (p-value < 0.01) across 241 gold standard genes was plotted using only missense variants from different allele frequency bins, where each bin includes all variants up to the specified maximum frequency. Scores were ranked from lowest to highest, and all thresholds were tested with a step size of 0.02. We selected the threshold per method with the highest Winsorized mean -log10(p-value) using singleton missense variants. The same threshold is used for all analyses in this figure. **B.** Winsorized Mean of -log10(p-value) Across 241 Gold Standard Genes. To account for outlier signals, we Winsorized the -log10(p-values) at 20.

Regeneron - Internal

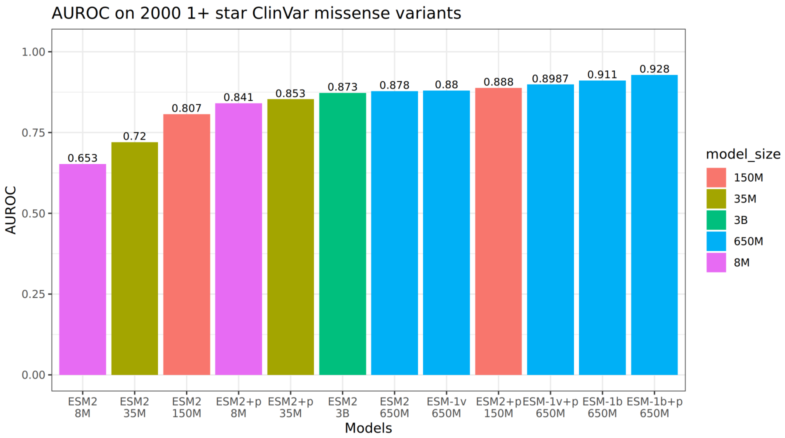


**Supplemental Figure 105: ESM model comparison on 2000 1+ plus star ClinVar variants.** We show the AUROC using ClinVar variants. The '+p' indicates proteomic refinement was used.
